# Investigating the effect of glycaemic traits on the risk of psychiatric illness using Mendelian randomisation

**DOI:** 10.1101/2020.03.09.984690

**Authors:** Danielle M. Adams, William R. Reay, Michael P. Geaghan, Murray J. Cairns

## Abstract

Data from observational studies have suggested an involvement of abnormal glycaemic regulation in the pathophysiology of psychiatric illness. This may be an attractive target for clinical intervention as glycaemia can be modulated by both lifestyle factors and pharmacological agents. However, observational studies are inherently confounded, and therefore causal relationships cannot be reliably established. We employed genetic variants rigorously associated with three glycaemic traits (fasting glucose, fasting insulin, and glycated haemoglobin) as instrumental variables in a two-sample Mendelian randomisation analysis to investigate the causal effect of these measures on the risk for eight psychiatric disorders. A significant protective effect of a unit increase in fasting insulin levels was observed for anorexia nervosa after the application of multiple testing correction (OR = 0.48 [95% CI: 0.33-0.71] – inverse-variance weighted estimate. The relationship between fasting insulin and anorexia nervosa was supported by a suite of sensitivity analyses, with no statistical evidence of instrument heterogeneity or horizontal pleiotropy. Further investigation is required to explore the relationship between insulin levels and anorexia.

## INTRODUCTION

Psychiatric disorders are complex phenotypes aetiologically influenced by a range of environmental (1, 2) and genetic factors (3–7). Currently, psychiatric disorders are treated with a combination of medication (8, 9) and psychotherapy approaches (10). Often these interventions address the symptoms of the disease without targeting the underlying mechanisms of action, and thus, managing psychiatric disorders remains difficult for many patients (11, 12). To address this, we need to better understand the risk factors and underlying pathophysiology of these conditions such that novel intervention strategies can be implemented.

There has been increasing interest in the relationship between glycaemic regulation and psychiatric illness. Dysglycaemia has well characterised systemic effects, however, its importance in the brain is often underappreciated. Insulin has been implicated in many neurological processes including synaptic plasticity and cognition (13–15), whilst neurons are dependent on glucose as their major energy source (16). A disproportionately high burden of comorbid type 2 diabetes has been observed in several psychiatric disorders, including: schizophrenia (17), bipolar disorder (17, 18), major depressive disorder (17), autism spectrum disorder (19), and Tourette's syndrome (20). In addition, glycaemic abnormalities have been observed through direct serum measurement, including an association of elevated glycated haemoglobin with attention deficit hyperactive disorder (21), insulin resistance with psychotic experiences (22), and increased insulin sensitivity in anorexia (23). Lifestyle factors and metabolic consequences of antipsychotic medication, such as weight gain (24, 25), likely contribute to these associations. However, data from treatment naïve, first episode psychosis patients provides evidence of glycaemic dysregulation in these disorders beyond what is directly attributable to medication effects and lifestyle (22, 26, 27). This relationship is further supported by genetic studies. For instance, linkage disequilibrium score regression (LDSC) has demonstrated a negative genetic correlation between anorexia and both fasting inulin and glucose as indexed by common genomic variation (28). Polygenic risk score for schizophrenia has also been associated with insulin resistance (29), whilst there is evidence of shared genome-wide association study (GWAS) association signals for schizophrenia and type 2 diabetes which display statistical colocalisation (30). Given the importance of glycaemic regulation in the brain and the direct significance of insulin signalling, these data suggest that dysglycaemia may be involved in the pathogenesis of psychiatric disorders. This could have implications for clinical monitoring and precision medicine as this system can be modulated through direct pharmacological intervention and lifestyle alterations.

The literature supporting the relationship between glycaemic traits and psychiatric disorders is largely composed of observational studies, preventing direct causal inferences. Randomised controlled trials (RCT) are viewed as an effective method to overcome this, however they are expensive and difficult to conduct with large sample sizes. An alternative method for inferring causal relationships between traits is Mendelian randomisation (MR), which is an analytical method to determine the causal effect of an exposure on an outcome by comparing the association of genetic instrumental variables (IV) with the outcome, relative to the IV effect on the exposure (31). Genetic variants which are rigorously associated with the exposure – discovered through GWAS – are selected as IVs, which in turn serve as proxies for the exposure. Two-sample MR is particularly advantageous as only GWAS summary statistics are required for the exposure and outcome traits of interest. Mendel's principle of independent assortment and random segregation underpin that these IVs will be randomized, thus their random distribution in the population emulates the random distribution of an exposure for individuals in a RCT (32, 33). In the present study, we have applied this approach to probe the causal effects of glycaemic traits on the risk for psychiatric disorders and observed a significant protective effect of increased fasting insulin levels on the risk of anorexia nervosa.

## MATERIALS AND METHODS

### Selection of genetic instrumental variables

Instrumental variables (IVs) for Mendelian randomisation are genetic variants associated with a particular effect size for a trait. There are three main assumptions which underlie the use of these instrumental variables (34–36):

IV1: the variant is rigorously associated with the exposure;

IV2: the variant is independent of all confounders of the exposure-outcome relationship (“exclusion-restriction assumption”); and

IV3: the variant is associated with the outcome only by acting through the exposure (independent conditional on the exposure and confounders).

IV1 is the only assumption which can be directly quantified (37); thus, we implement models (described below) to evaluate evidence for violations of these core assumptions. Specifically, pleiotropy, wherein a variant is associated with multiple phenotypes, may invalidate an IV if said pleiotropy constitutes an alternate causal pathway between the variant and the outcome (horizontal pleiotropy) (38).

We chose three core glycaemic traits to use as exposures in this study for which well-powered GWAS data were available: fasting insulin, fasting glucose, and glycated haemoglobin (HbA1c). IVs were genome-wide significant SNPs (*P* < 5 x 10^-8^, such that IV1 is satisfied) from the largest GWAS available for each trait (39, 40). Fasting insulin (FI) and fasting glucose (FG) data were obtained from the same meta-analysis of non-diabetic individuals of European ancestry (FI: N = 108,557, unit of effect = *ln* pmol/L; FG: N = 133,310, unit of effect = mmol/L). FI GWAS data was originally obtained from serum samples. FI results were additionally adjusted for body mass index (BMI) due to the complex relationship between insulin and weight gain (41–43). FG data was obtained from either plasma or from whole blood and corrected to plasma levels (39). IVs for HbA1c were obtained from the European subset of a GWAS meta-analysis (N = 123,665, unit of effect = % HbA1c). Genomewide significant SNPs were further categorised in this study as those acting through glycaemic pathways and those acting through erythrocytic pathways via annotation with GWAS catalog associations as described in Wheeler *et al.* (40). We chose to utilise the full set of significant SNPs as IVs, as well as the subset of the lead SNPs specifically annotated as glycaemic, to reduce potential horizontal pleiotropy (gHbAlc). The *F* statistic was calculated using equation one where *R^2^* is the variance in the outcome explained by each SNP, *k* is the number of viable IVs and *N* is the sample size ***(Table 1)***, demonstrating all IVs were sufficiently strong.

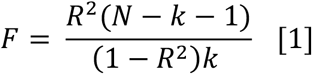

**Table 1:** Instrumental variables selected for each glycaemic exposure.

| Exposure | Number of IVs | Variance explained | F statistic | Sample size | Units |
| --- | --- | --- | --- | --- | --- |
| Fasting blood insulin | 14 | 0.64% | 47.59 | 108557 | $\ln$ pmol/L |
| Fasting glucose | 37 | 3.44% | 128.57 | 133310 | mmol/L |
| Fasting glycated haemoglobin (all) | 38 | 2.42% | 77.32 | 123665 | % glycated haemoglobin |
| Fasting glycated haemoglobin (glycaemic) | 15 | 0.85% | 67.50 | 123665 | % glycated haemoglobin |

The number of IVs, variance explained, $F$ statistic, sample size and units are described for the glycaemic exposures. The $F$ statistic was calculated from the number of IVs, variance explained and sample size as described previously (45). The variance explained was only from the IVs utilised in this study (44).

Hereafter, we refer to four exposures, as opposed to three, as there are two IV sets used for HbA1c, all SNPs and SNPs annotated as glycaemic (gHbA1c). IVs for all four exposures (FI, FG, HbA1c, and gHbA1c) were clumped to remove variants in linkage disequilibrium (r^2^ < 0.001) upon importation into the TwoSampleMR (version 0.4.25) R package (44) (R v3.6.1).

### Outcome data

We selected eight psychiatric disorders for which GWAS data were available as the outcome traits in this study: anorexia nervosa (AN), attention-deficit hyperactivity disorder (ADHD), autism spectrum disorder (ASD), bipolar disorder (BIP), major depressive disorder (MDD), obsessive compulsive disorder (OCD), schizophrenia (SZ), and Tourette's syndrome (TS). Outcome data was restricted to GWAS summary statistics from subjects of European ancestry in accordance with the exposure data, with the respective sample sizes as follows – AN: N = 72515 (46), ADHD: N = 53293 [European subset] (47), ASD: N = 46351 (48), BIP: N = 51710 (49), MDD: N = 1730005 [23andMe cohorts were not included in public release of the summary statistics from the psychiatric genomics consortium] (50), OCD: N = 9725 (51), SZ: N = 105318 (52), and TS: N = 14307 (53).

### Two sample Mendelian randomisation approach

An overview of the analysis workflow for this study is presented in ***Figure 1***. Firstly, we investigated the effect of fasting insulin, fasting glucose, HbA1c, and gHbA1c on the risk for each of the eight psychiatric disorders described above using an inverse-variance weighted effect model with multiplicative random effects (IVW) (54). The positive strand was inferred where possible otherwise palindromic SNPs were removed (55). We performed composite approaches and sensitivity analyses for exposure-outcome relationships which were significant after Bonferroni correction for the four exposures tested for eight outcomes [*P* < 1.56 × 10^-3^, *α* = 0.05/(8×4)].

**Figure 1:**
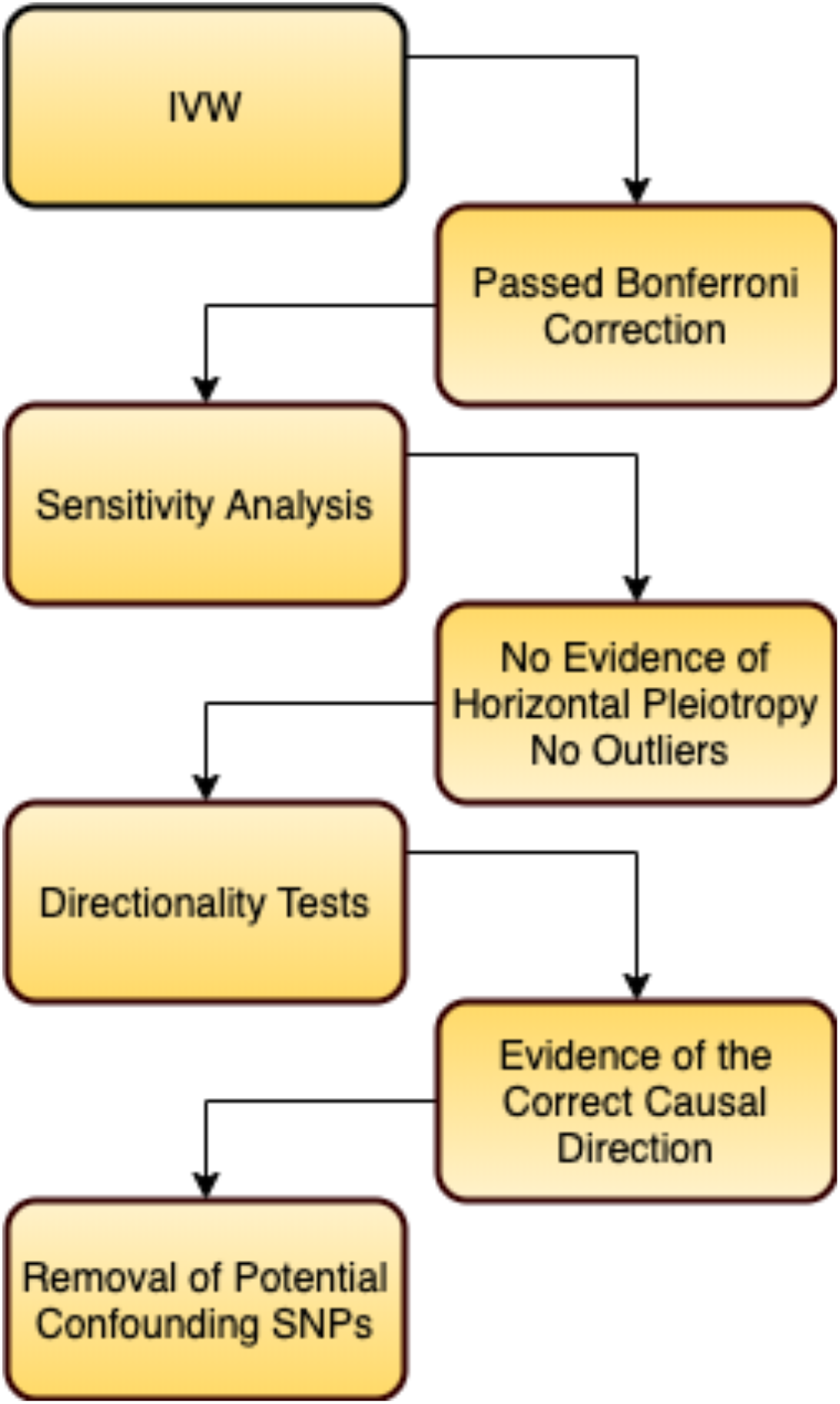
Workflow of the Mendelian randomisation analyses in this study. The effect of each glycaemic exposure on the eight psychiatric outcomes was tested using an inversevariance weighted effect model. Significant exposure-outcome causal estimates from this model, which survived multiple-testing correction using the Bonferroni method, were then retained for further sensitivity and pleiotropy analyses. Thereafter, evidence of horizontal pleiotropy and outlier instrumental variables was assessed. A directionality test was then implemented to ensure the exposure-outcome direction of causal effect was correct (as opposed to an outcome-exposure effect). IVW was then performed after the removal of potential confounding SNPs associated with both BMI and insulin.

The IVW model is limited such that even one invalid IV can bias the overall estimate. Therefore, for estimates with corrected significance we sought to overcome this limitation by using the outlier-robust MR-Pleiotropy Residual Sum and Outlier (MR-PRESSO) method (54, 56). MR-PRESSO is underpinned by the residual sum of squares (RSS), which serves as a heterogeneity measure of ratio estimates. Specifically, an IVW estimate using the IVs is calculated in a leave-one out fashion; if the RSS is decreased significantly relative to a simulated Gaussian distribution of expected RSS, then that variant is excluded from the IVW model. Simulations have demonstrated that this methodology is best suited to instances when less than half of the IVs exhibit horizontal pleiotropy (56). Two additional MR approaches were implemented to better account for potential invalid IVs: a weighted median estimate and MR-Egger. The weighted median model takes the median of the ratio estimates (as opposed to the mean in the IVW model), such that upweighting (with second order weights (57)) is applied to ratio estimates with greater precision (36). An advantage of this approach is that it is subject to the ‘majority valid' assumption, whereby an unbiased causal estimate will still be obtained if less than 50% of the model weighting arises from invalid IVs. Secondly, an MR-Egger model was constructed (35). This is an adaption of Egger regression wherein the exposure effect is regressed against the outcome with an intercept term added to represent the average pleiotropic effect.

### Sensitivity and pleiotropy analyses

The key assumption of the MR-Egger model is referred to as Instrument Strength Independent of Direct Effect (InSIDE), which assumes that there is no significant correlation between direct IV effects on the outcome and genetic association of IVs with the exposure (35, 58). In other words, the InSIDE assumption is violated if pleiotropic effects act through a confounder of the exposure-outcome association. We also tested whether the Egger intercept is significantly different from zero as a measure of unbalanced pleiotropy or violation of the InSIDE assumption (59). Furthermore, heterogeneity amongst the IV ratio estimates was quantified using Cochran's Q statistic, given that horizontal pleiotropy may be one explanation for significant heterogeneity (44, 60, 61). A global pleiotropy test was implemented via the MR-PRESSO framework utilising the expected and observed RSS (56). A leave-one-out analysis was then performed to assess whether causal estimates are biased by a single IV, which may indicate the presence of outliers, and the sensitivity of the estimate to said outliers (44). The MR Steiger directionality test utilizes the phenotypic variance explained by IV SNPs, comparing the instruments' association with the exposure and outcome to determine if there is evidence that the assumed direction of causality is correct (62). For binary traits, the trait population prevalence was used to calculate variance explained and convert to the liability scale using the lower and upper bounds of population prevalence estimates used by the GWAS for consistency (0.9% and 4% respectively for anorexia) (28, 63, 64). To investigate the significance of BMI-associated SNPs (65) on the relationship between outcome and exposure, SNPs associated with both traits were removed and the IVW estimate recalculated. All MR analyses were performed using the TwoSampleMR v0.4.25 package (44) in R v3.6.1 (66), with the exception of the MR-PRESSO model which utilised the MRPRESSO package v1.0.

### Latent causal variable model to estimate the genetic causality proportion of fasting insulin on risk for anorexia nervosa

Fasting insulin and anorexia nervosa display significant genome-wide genomic correlation as indexed by LDSC (46). This correlation may confound MR, and thus, we implemented a latent causal variable model (LCV) as an additional approach to investigate whether this correlation represents a causal relationship (67). Briefly, the LCV method assumes that a latent variable mediates the genetic correlation between two traits and tests whether this latent variable displays stronger correlation with either of the traits. Using fourth moments of the bivariate effect size distributions of all SNPs in both GWAS datasets and their LD structure, a posterior mean estimate of the genetic causality proportion (GCP) is derived which quantifies how much of the genomic architecture of one trait effects another. GCP values range from −1 to 1, with more positive values indicating greater partial genetic causality of trait one on two, and vice versa for more negative values. Full genetic causality is described as GCP = 1 or −1, which is rare in practice (67), with partial genetic causality occurring within these limits. A two-sided *t* test was used to assess whether the estimated GCP was significantly different from zero. The RunLCV.R and MomentFunctions.R scripts were leveraged to perform these analyses (https://github.com/lukejoconnor/LCV/tree/master/R). The fasting insulin summary statistics produced by Scott *et al.* were used as MR IVs followed up ~ 66,000 SNPs from previous GWAS. Whilst this is the largest sample size GWAS for this trait, the limited number of SNPs made it unsuitable for a genome-wide approach. Therefore, we utilised the smaller sample size GWAS summary statistics from Manning *et al.* (N = 33823) with more SNPs available as it was the basis for the LDSC previously performed between anorexia and fasting insulin (46, 68). Both summary statistics were cleaned and formatted in a standardised way (‘munged’) prior to analysis with the LCV model (67, 69, 70).

## RESULTS

Selected IVs explained approximately 3.44%, 0.64%, 2.42%, 0.85% of exposure variance of fasting glucose, insulin, and glycated haemoglobin levels (HbA1c, gHbA1c) respectively. IVs were selected by clumping for LD to remove correlated variants and excluding palindromes for which the correct strand could not be inferred. An IVW model was used to estimate the casual effect of the four exposures on the eight psychiatric disorder outcomes. We revealed a significant protective effect of a unit increase [*ln*(pmol/L)] in fasting insulin levels on anorexia after the applying Bonferroni correction [OR=0.48, 95% CI:0.33-0.71, *P*=2.27 × 10^-4^] ***(Figure 2,3)***. In addition, the relationship between fasting insulin and MDD was nominally significant (uncorrected *P* < 0.05) [OR=0.85, 95% CI: 0.74-0.97, *P*=0.015], whilst all other causal estimates were not significant ***(Supplementary table 1)*.**

**Figure 2:**
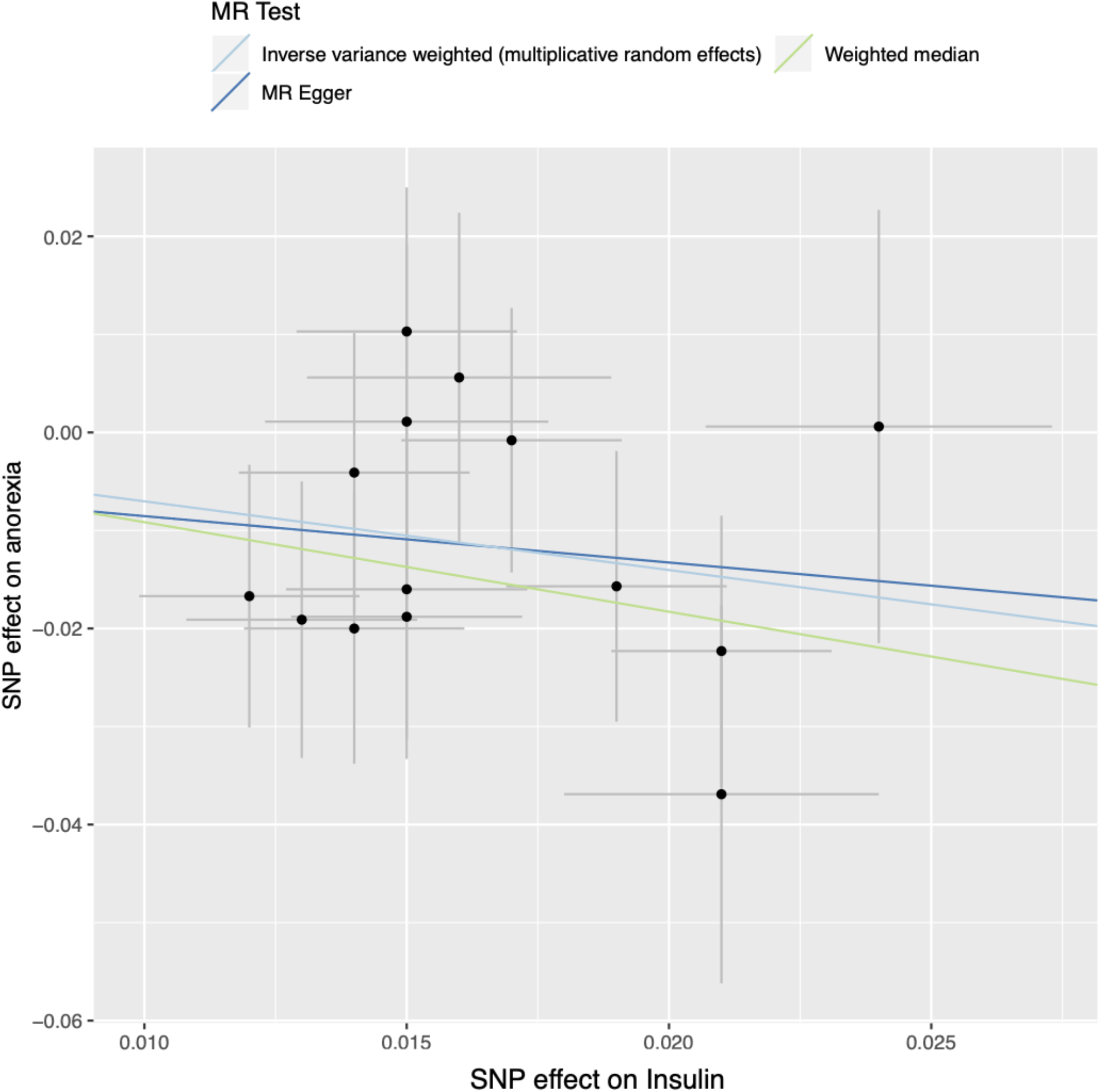
Effects of genetic instrumental variables on fasting insulin and anorexia nervosa. Points represent individual single nucleotide polymorphisms (SNPs) with grey lines indicating 95% confidence intervals. The legend denotes which MR test was used; inverse variance weighted (multiplicative random effects), MR egger or the weighted median.

**Figure 3:**
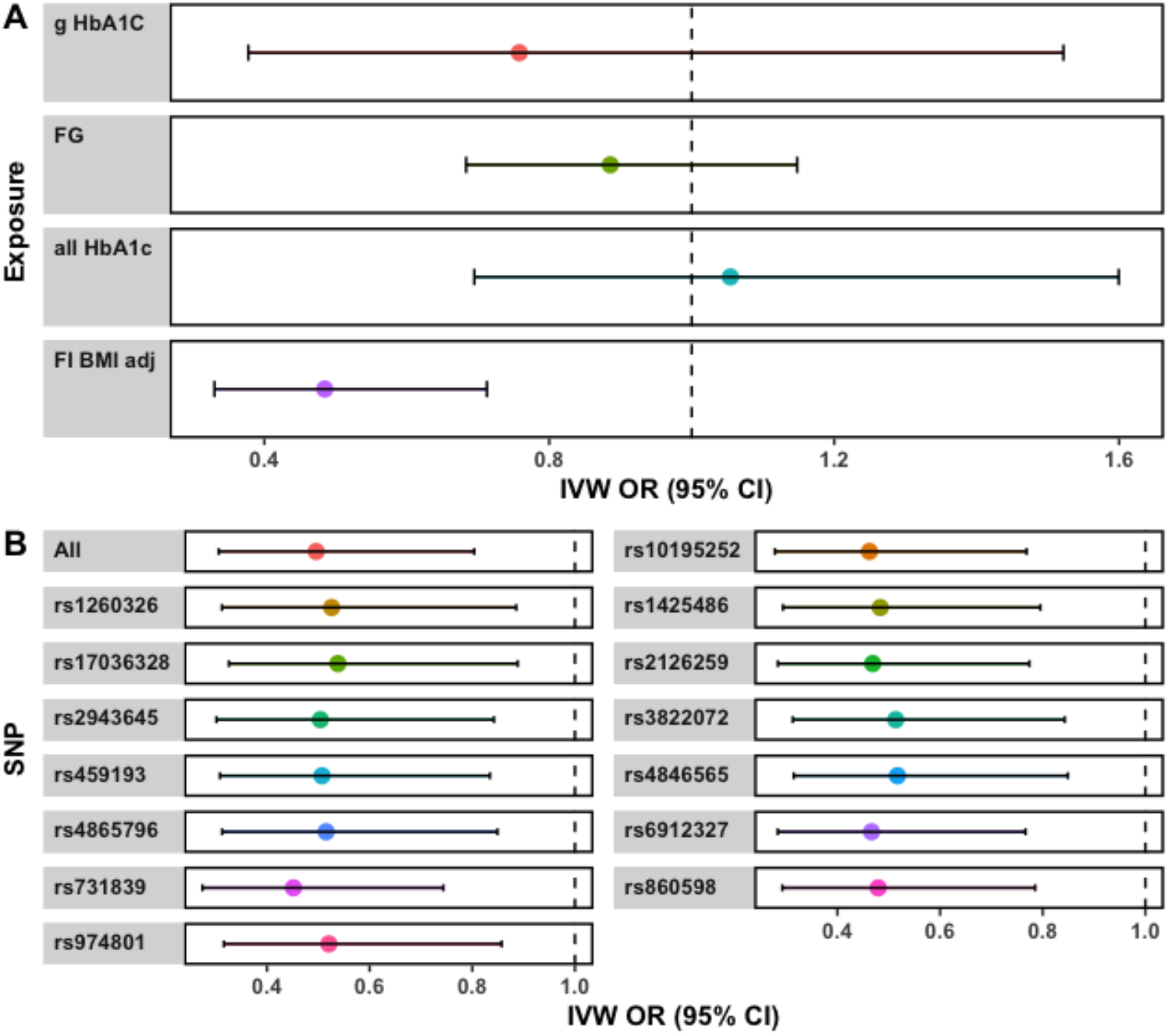
Increasing fasting insulin is associated with a protective effect on risk for anorexia nervosa. **A:** Effect size estimates for the glycaemic exposures on anorexia nervosa. The inverse variance weighted effect model was utilised to calculate point estimates for each exposure as visualised on the Forest plot, with 95% confidence intervals shown. FI BMI adj=fasting insulin BMI adjusted, FG= fasting glucose, all HbA1c= all glycated haemoglobin SNPs, gHbA1c= glycaemic annotated glycated haemoglobin SNPs. **B:** Leave one out recalculation of the effect of fasting insulin on anorexia nervosa. The vertical axis shows which SNP was removed and the causal estimate recalculated; ‘All’ indicates the effect estimate when no SNPs were removed.

We subjected the causal estimate of fasting insulin on the risk for anorexia to a suite of sensitivity analyses to assess the rigor of our derived IVW estimate and evidence of violations of core MR assumptions. No IVs were detected as outliers using the MR-PRESSO approach. Furthermore, the weighted median model supported the putative protective effect of increasing fasting insulin on risk for anorexia derived from the IVW (OR= 0.40, 95% CI:0.21-0.77, *P* = 6.3 × 10^-3^). The MR-Egger model was not significant; however, the causal estimate was in the same direction of effect as the other two approaches, albeit with an extremely wide confidence interval (OR = 0.68, 95% CI: 0.05-9.10). It should be noted that the MR-Egger method typically has notably less power than other approaches, particularly when fewer IVs are used (35).There was no compelling evidence for unbalanced pleiotropy amongst IVs utilised in this construct: heterogeneity between IV effects was not significant (*Q* = 8.24, *df* = 13, *P* = 0.827), the intercept of the MR egger regression did not significantly differ from zero (intercept = −5.7 × 10^-3^, *P* = 0.795), and the MR PRESSO test of global pleiotropy was also not significant. A leave-one out recalculation of the causal estimate did not indicate that a single IV or subset of IVs were unduly influencing the model ***(Figure 3)***. Given the putative bidirectional relationship between anorexia and body mass index (BMI) (46), we identified five fasting insulin IVs which were also associated with BMI at genome wide significance (*P* < 5 × 10^-8^) and recalculated the IVW estimate with these instruments removed. The effect size observed in this reduced IVW model was not greatly attenuated (OR = 0.51 [95% CI: 0.32 – 0.83], *P* = 7.03 × 10^-3^), suggesting that the relationship between insulin and anorexia is not unduly biased by horizontal pleiotropy through IV effects on BMI.

The MR models implemented in this study assume that the IVs impact the exposure, which in turn affects the outcome – however, in practice it is feasible that the orientation of the causal pathway is incorrect and that the outcome influences the exposure through the genetic instruments. To address this, we performed a Steiger directionality test. The variance in anorexia risk explained by the IVs was converted to the liability scale using the upper (4%) and lower (0.9%) bound of estimated population prevalence for anorexia (28), which supported the hypothesis that the effect of fasting insulin on risk for anorexia is the correct causal direction (*P* = 1.35 × 10^-27^, *P* = 1.36 × 10^-27^ respectively for the upper and lower bound prevalence estimates) ***(Supplementary table 5)***. Given the genetic correlation between fasting insulin and anorexia, we used a latent causal variable model to determine the proportion of trait one (fasting insulin) that genetically causes trait two (anorexia), which was quantified as the mean posterior estimate of the genetic causality proportion (GCP). The sign of the mean posterior GCP estimate suggests that fasting insulin is partially genetically causal for anorexia; however, this was not significantly different from zero, likely due to the large standard error 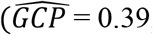, *SE* = 0.33, *P*=0.26]. Whilst the LCV GCP estimate was not significant, there was strong evidence that the causal direction was not anorexia to insulin [H_0_: GCP= −1, P=3.6 × 10^-104^], in accordance with the Steiger directionality test results.

## DISCUSSION

Glycaemic regulation is involved with many physiological processes, however, its role in neurological function has motivated investigation of this system in psychiatric disorders. Using a Mendelian randomisation approach which leverages genetic IVs as proxies for three glycaemic traits, we uncovered evidence of a protective effect of increasing fasting insulin on the risk for anorexia nervosa. No significant evidence of a causal effect of any of the glycaemic traits was found testing seven other psychiatric phenotypes. Notably, we did not replicate a previous study which demonstrated a risk increasing effect of fasting insulin on schizophrenia, however, we utilised a larger schizophrenia GWAS and different IVs (71). Although previous analysis has shown a relationship between first episode psychosis and glycaemic dysregulation, and elevated rates of dysglycaemia in psychiatry, this was not supported by our MR model. Our inability to detect this relationship may be limited by the strength of the instrumental variables used, or the observed effects from previous analysis may be due to variables with shared genetic liability which influence glycaemic homeostasis, such as inflammation or BMI. The future availability of more data with the power to explain a larger portion of the variance in the exposures and outcomes, could yet yield more evidence of a causal relationship between dysglycaemia and other psychiatric disorders. The negative relationship between increasing insulin and anorexia risk derived in this study supports the negative genetic correlation observed between the two GWAS studies by LDSC (28). The association between a natural log transformed pmol/L increase in fasting insulin and odds of anorexia yielded an odds ratio of 0.48 [95% CI: 0.33-0.71]. To contextualize this unit of effect, we considered fasting insulin values from a large cohort of 10.5 to 11 year old female normal weight European participants (72). The bottom decile of this cohort was estimated to have a 30.97 pmol/L fasting insulin concentration, a unit increase to which would correspond to approximately 84.19 pmol/L, roughly equivalent to the 90^th^ percentile of the cohort (~ 86.91). This estimate derived from the IVW model was supported by sensitivity analyses which did not indicate any statistical evidence of unbalanced pleiotropy which would confound the IVs we selected.

The role of insulin signalling and glucose metabolism in the brain, and its interplay with the periphery, is complex, necessitating further research to specifically understand how fasting insulin could exert a protective effect on anorexia. The relationship between circulating insulin and weight gain may contribute to this protective effect given peripheral insulin and insulin therapy in the context of diabetes is associated with weight gain (73, 74). There are a number of mechanisms by which this is proposed to be mediated, including the stimulatory effect of insulin on fatty acid storage and cell growth and a reduction in glycosuria (75–77). Furthermore, Mendelian randomisation analyses have supported a positive relationship between insulin and weight gain (78, 79). Given the nature of the clinical presentation of anorexia, the effect of insulin on hunger and satiety is particularly pertinent. Increased insulin levels in the body results in higher levels of hunger and an increased pleasantness associated with sweet taste (80). This corresponds to data from anorexia cohorts which report that individuals with anorexia have a reduced appreciation of sweet tastes (81). In contrast, insulin is postulated to have an anorexigenic effect in the brain (82, 83), partly through its inhibition of the orexigenic agouti-related peptide (AgRP) and neuropeptide Y (NPY) neurons (84, 85). This may contradict the putative risk-decreasing effect of insulin on anorexia we observed in our study, however, there is evidence of a significant sexual dimorphism in this phenomenon. An example of this has been demonstrated using intranasal insulin administration, in which hunger was decreased only in male participants, conversely, there were positive cognitive enhancing effects seen only in women (86). This sexual dimorphism in the effect of insulin signalling is further supported by rodent data (87). As anorexia is significantly more prevalent in females (88) it is possible that the sexual dimorphic effect of insulin on hunger signalling is contributing to this discrepancy in prevalence.

There are a number of future directions which arise from these data. Only Europeans were used in this analysis warranting its extension to trans-ethnic cohorts. As the negative relationship between insulin and anorexia is also evidenced using a genomic correlation approach, there is a need to investigate shared genes and biological pathways which may explain this association. This would be particularly valuable to interpret the causal estimate we uncovered, as individuals who develop anorexia may be genetically predisposed to have altered glycaemic homeostasis. Furthermore, a well-powered, sex stratified GWAS of anorexia could be utilised to formally test whether the impact of insulin on anorexia risk displays sexual dimorphism. As diagnosis of anorexia is highly skewed towards females, it will likely be a continued challenge to genotype larger male cohorts which approach the sample sizes available for female participants. It is also important to consider the inherent limitations of MR in light of our data. We did not uncover any statistical evidence of unbalanced pleiotropy amongst the fasting insulin IVs; however, this cannot be definitively proven and future replication in larger studies is paramount. Moreover, whilst the IVs selected for insulin were appropriately strong as quantified by an *F*-statistic, they still only explain a fraction of the phenotypic variance in fasting insulin. Despite these caveats and other methodological challenges associated with causal inference using IVs, we believe the insulin – anorexia model to be reliable. In conclusion, we uncovered evidence of a protective effect of fasting insulin on the risk of anorexia nervosa, with further work now required to further understand the biological mechanisms underpinning this relationship.

## REFERENCES

1. McGrath JJ, Saha S, Lim CCW, Aguilar-Gaxiola S, Alonso J, Andrade LH, et al. Trauma and psychotic experiences: transnational data from the World Mental Health Survey. Br J Psychiatry. 2017;211(6):373–80.

2. van Os J, Kenis G, Rutten BPF. The environment and schizophrenia. Nature. 2010;468(7321):203–12.

3. Cross-Disorder Group of the Psychiatric Genomics Consortium. Electronic address pmhe, Cross-Disorder Group of the Psychiatric Genomics C. Genomic Relationships, Novel Loci, and Pleiotropic Mechanisms across Eight Psychiatric Disorders. Cell. 2019;179(7):1469–82.e11.

4. Polderman TJC, Benyamin B, de Leeuw CA, Sullivan PF, van Bochoven A, Visscher PM, et al. Meta-analysis of the heritability of human traits based on fifty years of twin studies. Nat Genet. 2015;47(7):702–9.

5. Duncan LE, Ostacher M, Ballon J. How genome-wide association studies (GWAS) made traditional candidate gene studies obsolete. Neuropsychopharmacology. 2019;44(9):1518–23.

6. Nestadt G, Grados M, Samuels JF. Genetics of obsessive-compulsive disorder. The Psychiatric clinics of North America. 2010;33(1):141–58.

7. Sullivan PF, Daly MJ, O'Donovan M. Genetic architectures of psychiatric disorders: the emerging picture and its implications. Nat Rev Genet. 2012;13(8):537–51.

8. Henderson DC. Managing weight gain and metabolic issues in patients treated with atypical antipsychotics. J Clin Psychiatry. 2008;69(2):e04.

9. Lake J, Turner MS. Urgent Need for Improved Mental Health Care and a More Collaborative Model of Care. Perm J. 2017;21:17–024.

10. Hofmann SG, Asnaani A, Vonk IJJ, Sawyer AT, Fang A. The Efficacy of Cognitive Behavioral Therapy: A Review of Meta-analyses.Cognit Ther Res. 2012;36(5):427–40.

11. Howes OD, McCutcheon R, Agid O, de Bartolomeis A, van Beveren NJM, Birnbaum ML, et al. Treatment-Resistant Schizophrenia: Treatment Response and Resistance in Psychosis (TRRIP) Working Group Consensus Guidelines on Diagnosis and Terminology. The American journal of psychiatry. 2017;174(3):216–29.

12. Rush AJ, Trivedi MH, Wisniewski SR, Nierenberg AA, Stewart JW, Warden D, et al. Acute and longer-term outcomes in depressed outpatients requiring one or several treatment steps: a STAR*D report. The American journal of psychiatry. 2006;163(11):1905–17.

13. Biessels GJ, Kamal A, Urban IJ, Spruijt BM, Erkelens DW, Gispen WH. Water maze learning and hippocampal synaptic plasticity in streptozotocin-diabetic rats: effects of insulin treatment. Brain research. 1998;800(1):125–35.

14. Grillo CA, Piroli GG, Lawrence RC, Wrighten SA, Green AJ, Wilson SP, et al. Hippocampal Insulin Resistance Impairs Spatial Learning and Synaptic Plasticity. Diabetes. 2015;64(11):3927–36.

15. Duarte AI, Moreira PI, Oliveira CR. Insulin in Central Nervous System: More than Just a Peripheral Hormone. Journal of Aging Research. 2012;2012:21.

16. Mergenthaler P, Lindauer U, Dienel GA, Meisel A. Sugar for the brain: the role of glucose in physiological and pathological brain function. Trends Neurosci. 2013;36(10):587–97.

17. Vancampfort D, Correll CU, Galling B, Probst M, De Hert M, Ward PB, et al. Diabetes mellitus in people with schizophrenia, bipolar disorder and major depressive disorder: a systematic review and large scale meta-analysis. World Psychiatry. 2016;15(2):166–74.

18. Vancampfort D, Mitchell AJ, De Hert M, Sienaert P, Probst M, Buys R, et al. Prevalence and predictors of type 2 diabetes mellitus in people with bipolar disorder: a systematic review and meta-analysis. J Clin Psychiatry. 2015;76(11):1490–9.

19. Chen MH, Lan WH, Hsu JW, Huang KL, Su TP, Li CT, et al. Risk of Developing Type 2 Diabetes in Adolescents and Young Adults With Autism Spectrum Disorder: A Nationwide Longitudinal Study. Diabetes Care. 2016;39(5):788–93.

20. Brander G, Isomura K, Chang Z, Kuja-Halkola R, Almqvist C, Larsson H, et al. Association of Tourette Syndrome and Chronic Tic Disorder With Metabolic and Cardiovascular Disorders. JAMA Neurol. 2019;76(4):454–61.

21. Lindblad F, Eickhoff M, Forslund AH, Isaksson J, Gustafsson J. Fasting blood glucose and HbA1c in children with ADHD. Psychiatry Res. 2015;226(2-3):515–6.

22. Perry BI, Upthegrove R, Thompson A, Marwaha S, Zammit S, Singh SP, et al. Dysglycaemia, Inflammation and Psychosis: Findings From the UK ALSPAC Birth Cohort. Schizophr Bull. 2019;45(2):330–8.

23. Dostalova I, Smitka K, Papezova H, Kvasnickova H, Nedvidkova J. Increased insulin sensitivity in patients with anorexia nervosa: the role of adipocytokines. Physiol Res. 2007;56(5):587–94.

24. Schimmelmann BG, Schmidt SJ, Carbon M, Correll CU. Treatment of adolescents with early-onset schizophrenia spectrum disorders: in search of a rational, evidence-informed approach. Curr Opin Psychiatry. 2013;26(2):219–30.

25. Correll CU, Detraux J, De Lepeleire J, De Hert M. Effects of antipsychotics, antidepressants and mood stabilizers on risk for physical diseases in people with schizophrenia, depression and bipolar disorder. World Psychiatry. 2015;14(2):119–36.

26. Pillinger T, Beck K, Gobjila C, Donocik JG, Jauhar S, Howes OD. Impaired Glucose Homeostasis in First-Episode Schizophrenia: A Systematic Review and Meta-analysis. JAMA Psychiatry. 2017;74(3):261–9.

27. Perry BI, McIntosh G, Weich S, Singh S, Rees K. The association between first-episode psychosis and abnormal glycaemic control: systematic review and meta-analysis. Lancet Psychiatry. 2016;3(11):1049–58.

28. Watson H, Yilmaz Z, Thornton L, Hübel C, Coleman J, Gaspar H, et al. Genome-wide association study identifies eight risk loci and implicates metabo-psychiatric origins for anorexia nervosa. Nat Genet. 2019;51.

29. Tomasik J, Lago SG, Vázquez-Bourgon J, Papiol S, Suárez-Pinilla P, Crespo-Facorro B, et al. Association of Insulin Resistance With Schizophrenia Polygenic Risk Score and Response to Antipsychotic TreatmentInsulin Resistance and Schizophrenia Polygenic Risk Score and Response to Antipsychotic TreatmentLetters. JAMA Psychiatry. 2019;76(8):864–7.

30. Hackinger S, Prins B, Mamakou V, Zengini E, Marouli E, Brčić L, et al. Evidence for genetic contribution to the increased risk of type 2 diabetes in schizophrenia. Translational Psychiatry. 2018;8(1):252.

31. Rasooly D, Patel CJ. Conducting a Reproducible Mendelian Randomization Analysis Using the R Analytic Statistical Environment. Current Protocols in Human Genetics. 2019;101(1):e82.

32. Lawlor DA, Harbord RM, Sterne JAC, Timpson N, Davey Smith G. Mendelian randomization: Using genes as instruments for making causal inferences in epidemiology. Statistics in Medicine. 2008;27(8):1133–63.

33. Teumer A. Common Methods for Performing Mendelian Randomization. Frontiers in cardiovascular medicine. 2018;5:51.

34. Martens EP, Pestman WR, de Boer A, Belitser SV, Klungel OH. Instrumental variables: application and limitations. Epidemiology (Cambridge, Mass). 2006;17(3):260–7.

35. Bowden J, Davey Smith G, Burgess S. Mendelian randomization with invalid instruments: effect estimation and bias detection through Egger regression. International Journal of Epidemiology. 2015;44(2):512–25.

36. Bowden J, Davey Smith G, Haycock PC, Burgess S. Consistent Estimation in Mendelian Randomization with Some Invalid Instruments Using a Weighted Median Estimator. Genetic epidemiology. 2016;40(4):304–14.

37. Labrecque J, Swanson SA. Understanding the Assumptions Underlying Instrumental Variable Analyses: a Brief Review of Falsification Strategies and Related Tools. Current Epidemiology Reports. 2018;5(3):214–20.

38. Lawlor DA, Harbord RM, Sterne JA, Timpson N, Davey Smith G. Mendelian randomization: using genes as instruments for making causal inferences in epidemiology. Statistics in medicine. 2008;27(8):1133–63.

39. Scott RA, Lagou V, Welch RP, Wheeler E, Montasser ME, Luan Ja, et al. Large-scale association analyses identify new loci influencing glycemic traits and provide insight into the underlying biological pathways. Nat Genet. 2012;44(9):991–1005.

40. Wheeler E, Leong A, Liu C-T, Hivert M-F, Strawbridge RJ, Podmore C, et al. Impact of common genetic determinants of Hemoglobin A1c on type 2 diabetes risk and diagnosis in ancestrally diverse populations: A transethnic genome-wide meta-analysis. PLoS medicine. 2017;14(9):e1002383.

41. Scott RA, Fall T, Pasko D, Barker A, Sharp SJ, Arriola L, et al. Common genetic variants highlight the role of insulin resistance and body fat distribution in type 2 diabetes, independent of obesity. Diabetes. 2014;63(12):4378–87.

42. Gonzalez-Cantero J, Martin-Rodriguez JL, Gonzalez-Cantero A, Arrebola JP, Gonzalez-Calvin JL. Insulin resistance in lean and overweight non-diabetic Caucasian adults: Study of its relationship with liver triglyceride content, waist circumference and BMI. PLOS ONE. 2018;13(2):e0192663.

43. Pennings N, Jaber J, Ahiawodzi P. Ten-year weight gain is associated with elevated fasting insulin levels and precedes glucose elevation. Diabetes/metabolism research and reviews. 2018;34(4):e2986.

44. Hemani G, Zheng J, Elsworth B, Wade KH, Haberland V, Baird D, et al. The MR-Base platform supports systematic causal inference across the human phenome. eLife. 2018;7:e34408.

45. Shu X, Wu L, Khankari NK, Shu X-O, Wang TJ, Michailidou K, et al. Associations of obesity and circulating insulin and glucose with breast cancer risk: a Mendelian randomization analysis. International Journal of Epidemiology. 2018;48(3):795–806.

46. Watson HJ, Yilmaz Z, Thornton LM, Hübel C, Coleman JRI, Gaspar HA, et al. Genomewide association study identifies eight risk loci and implicates metabo-psychiatric origins for anorexia nervosa. Nature genetics. 2019;51(8):1207–14.

47. Demontis D, Walters RK, Martin J, Mattheisen M, Als TD, Agerbo E, et al. Discovery of the first genome-wide significant risk loci for attention deficit/hyperactivity disorder. Nature genetics. 2019;51(1):63–75.

48. Grove J, Ripke S, Als TD, Mattheisen M, Walters RK, Won H, et al. Identification of common genetic risk variants for autism spectrum disorder. Nature genetics. 2019;51(3):431–44.

49. Stahl EA, Breen G, Forstner AJ, McQuillin A, Ripke S, Trubetskoy V, et al. Genome-wide association study identifies 30 loci associated with bipolar disorder. Nature genetics. 2019;51(5):793–803.

50. Wray NR, Ripke S, Mattheisen M, Trzaskowski M, Byrne EM, Abdellaoui A, et al. Genome-wide association analyses identify 44 risk variants and refine the genetic architecture of major depression. Nature genetics. 2018;50(5):668–81.

51. OCD IOCDFGCI-Ga, (OCGAS). CGAS. Revealing the complex genetic architecture of obsessive-compulsive disorder using meta-analysis. Mol Psychiatry. 2018;23(5):1181–8.

52. Pardiñas AF, Holmans P, Pocklington AJ, Escott-Price V, Ripke S, Carrera N, et al. Common schizophrenia alleles are enriched in mutation-intolerant genes and in regions under strong background selection. Nature genetics. 2018;50(3):381–9.

53. Yu D, Sul JH, Tsetsos F, Nawaz MS, Huang AY, Zelaya I, et al. Interrogating the Genetic Determinants of Tourette's Syndrome and Other Tic Disorders Through Genome-Wide Association Studies. The American journal of psychiatry. 2019;176(3):217–27.

54. Burgess S, Butterworth A, Thompson SG. Mendelian randomization analysis with multiple genetic variants using summarized data. Genetic epidemiology. 2013;37(7):658–65.

55. Hartwig FP, Davies NM, Hemani G, Davey Smith G. Two-sample Mendelian randomization: avoiding the downsides of a powerful, widely applicable but potentially fallible technique. Int J Epidemiol. 2016;45(6):1717–26.

56. Verbanck M, Chen C-Y, Neale B, Do R. Detection of widespread horizontal pleiotropy in causal relationships inferred from Mendelian randomization between complex traits and diseases. Nature genetics. 2018;50(5):693–8.

57. Bowden J, Del Greco M F, Minelli C, Zhao Q, Lawlor DA, Sheehan NA, et al. Improving the accuracy of two-sample summary-data Mendelian randomization: moving beyond the NOME assumption. Int J Epidemiol. 2018;48(3):728–42.

58. Slob EAW, Burgess S. A Comparison Of Robust Mendelian Randomization Methods Using Summary Data. bioRxiv. 2019:577940.

59. Burgess S, Thompson SG. Interpreting findings from Mendelian randomization using the MR-Egger method. Eur J Epidemiol. 2017;32(5):377–89.

60. Cochran WG. THE COMPARISON OF PERCENTAGES IN MATCHED SAMPLES. Biometrika. 1950;37(3-4):256–66.

61. Bowden J, Hemani G, Davey Smith G. Invited Commentary: Detecting Individual and Global Horizontal Pleiotropy in Mendelian Randomization—A Job for the Humble Heterogeneity Statistic? American Journal of Epidemiology. 2018;187(12):2681–5.

62. Hemani G, Tilling K, Davey Smith G. Orienting the causal relationship between imprecisely measured traits using GWAS summary data. PLoS Genet. 2017;13(11):e1007081–e.

63. Lee SH, Wray NR. Novel genetic analysis for case-control genome-wide association studies: quantification of power and genomic prediction accuracy. PLoS One. 2013;8(8):e71494–e.

64. Lee SH, Goddard ME, Wray NR, Visscher PM. A Better Coefficient of Determination for Genetic Profile Analysis. Genetic Epidemiology. 2012;36(3):214–24.

65. Pulit SL, Stoneman C, Morris AP, Wood AR, Glastonbury CA, Tyrrell J, et al. Metaanalysis of genome-wide association studies for body fat distribution in 694 649 individuals of European ancestry. Human molecular genetics. 2019;28(1):166–74.

66. Team R. A language and environment for statistical computing. Computing. 2006;1.

67. O'Connor LJ, Price AL. Distinguishing genetic correlation from causation across 52 diseases and complex traits. Nature genetics. 2018;50(12):1728–34.

68. Manning AK, Hivert M-F, Scott RA, Grimsby JL, Bouatia-Naji N, Chen H, et al. A genome-wide approach accounting for body mass index identifies genetic variants influencing fasting glycemic traits and insulin resistance. Nature genetics. 2012;44(6):659–69.

69. Anttila V, Bulik-Sullivan B, Finucane HK, Walters RK, Bras J, Duncan L, et al. Analysis of shared heritability in common disorders of the brain. Science (New York, NY). 2018;360(6395).

70. Bulik-Sullivan BK, Loh P-R, Finucane HK, Ripke S, Yang J, Patterson N, et al. LD Score regression distinguishes confounding from polygenicity in genome-wide association studies. Nature genetics. 2015;47(3):291–5.

71. Li Z, Chen P, Chen J, Xu Y, Wang Q, Li X, et al. Glucose and Insulin-Related Traits, Type 2 Diabetes and Risk of Schizophrenia: A Mendelian Randomization Study. EBioMedicine. 2018;34:182–8.

72. Peplies J, Jiménez-Pavón D, Savva SC, Buck C, Günther K, Fraterman A, et al. Percentiles of fasting serum insulin, glucose, HbA1c and HOMA-IR in pre-pubertal normal weight European children from the IDEFICS cohort. International Journal of Obesity. 2014;38(2):S39–S47.

73. Pennings N, Jaber J, Ahiawodzi P. Ten-year weight gain is associated with elevated fasting insulin levels and precedes glucose elevation. Diabetes Metab Res Rev. 2018;34(4):e2986–e.

74. Brown A, Guess N, Dornhorst A, Taheri S, Frost G. Insulin-associated weight gain in obese type 2 diabetes mellitus patients: What can be done? Diabetes Obes Metab. 2017;19(12):1655–68.

75. Packianathan IC, Fuller NJ, Peterson DB, Wright A, Coward WA, Finer N. Use of a reference four-component model to define the effects of insulin treatment on body composition in type 2 diabetes: the 'Darwin study'. Diabetologia. 2005;48(2):222–9.

76. Mäkimattila S, Nikkilä K, Yki-Järvinen H. Causes of weight gain during insulin therapy with and without metformin in patients with Type II diabetes mellitus. Diabetologia. 1999;42(4):406–12.

77. Shank ML, Del Prato S, DeFronzo RA. Bedtime insulin/daytime glipizide. Effective therapy for sulfonylurea failures in NIDDM. Diabetes. 1995;44(2):165–72.

78. Xu L, Borges MC, Hemani G, Lawlor DA. The role of glycaemic and lipid risk factors in mediating the effect of BMI on coronary heart disease: a two-step, two-sample Mendelian randomisation study. Diabetologia. 2017;60(11):2210–20.

79. Astley CM, Todd JN, Salem RM, Vedantam S, Ebbeling CB, Huang PL, et al. Genetic Evidence That Carbohydrate-Stimulated Insulin Secretion Leads to Obesity. Clin Chem. 2018;64(1):192–200.

80. Rodin J. Insulin levels, hunger, and food intake: an example of feedback loops in body weight regulation. Health Psychol. 1985;4(1):1–24.

81. Kinnaird E, Stewart C, Tchanturia K. Taste sensitivity in anorexia nervosa: A systematic review. Int J Eat Disord. 2018;51(8):771–84.

82. Kullmann S, Heni M, Veit R, Scheffler K, Machann J, Häring H-U, et al. Intranasal insulin enhances brain functional connectivity mediating the relationship between adiposity and subjective feeling of hunger. Scientific Reports. 2017;7(1):1627.

83. Plum L, Schubert M, Brüning JC. The role of insulin receptor signaling in the brain. Trends in Endocrinology & Metabolism. 2005;16(2):59–65.

84. Loh K, Zhang L, Brandon A, Wang Q, Begg D, Qi Y, et al. Insulin controls food intake and energy balance via NPY neurons. Mol Metab. 2017;6(6):574–84.

85. Kaga T, Inui A, Okita M, Asakawa A, Ueno N, Kasuga M, et al. Modest Overexpression of Neuropeptide Y in the Brain Leads to Obesity After High-Sucrose Feeding. Diabetes. 2001;50(5):1206–10.

86. Benedict C, Kern W, Schultes B, Born J, Hallschmid M. Differential sensitivity of men and women to anorexigenic and memory-improving effects of intranasal insulin. J Clin Endocrinol Metab. 2008;93(4):1339–44.

87. Clegg DJ, Riedy CA, Smith KAB, Benoit SC, Woods SC. Differential Sensitivity to Central Leptin and Insulin in Male and Female Rats. Diabetes. 2003;52(3):682.

88. Klump KL. Puberty as a critical risk period for eating disorders: a review of human and animal studies. Horm Behav. 2013;64(2):399–410.

